# LUC7 proteins define two major classes of 5’ splice sites in animals and plants

**DOI:** 10.1101/2022.12.07.519539

**Authors:** Connor J. Kenny, Michael P. McGurk, Sandra Schüler, Aidan Cordero, Sascha Laubinger, Christopher B. Burge

## Abstract

Mutation or deletion of the U1 snRNP-associated factor LUC7L2 is associated with myeloid neoplasms, and knockout of LUC7L2 alters cellular metabolism. Here, we uncover that members of the LUC7 protein family differentially regulate two major classes of 5’ splice sites (5’SS) and broadly regulate mRNA splicing in both human cell lines and leukemias with *LUC7L2* copy number variation. We describe distinctive 5’SS features of exons impacted by the three human LUC7 paralogs: LUC7L2 and LUC7L enhance splicing of “right-handed” 5’SS with stronger consensus matching on the intron side of the near-invariant /GU, while LUC7L3 enhances splicing of “left-handed” 5’SS with stronger consensus matching upstream of the /GU. We validated our model of sequence-specific 5’SS regulation both by mutating splice sites and swapping domains between human LUC7 proteins. Evolutionary analysis indicates that the LUC7L2/LUC7L3 subfamilies diverged before the divergence of animals and plants. Analysis of *Arabidopsis thaliana* mutants confirmed that plant LUC7 orthologs possess specificity similar to their human counterparts, indicating that 5’SS regulation by LUC7 proteins is deeply conserved.

## Introduction

Eukaryotic protein-coding genes are often interrupted by non-coding introns, which are recognized and excised by the spliceosome. In the earliest steps of spliceosome assembly, U1 snRNP identifies potential intron boundaries in pre-mRNAs via RNA:RNA base pairing between the 5’ end of U1 snRNA and the 5’ splice site (5SS), a conserved 9 nucleotide motif that demarcates the exon-intron boundary (Krämer et al., 1984). While the 5’ end of U1 snRNA is nearly invariant, there is considerable variation in functional 5’SS motifs between eukaryotes (Figure 1A, Schwartz et al., 2008, Wong et al., 2016) indicating additional factors contribute to the recognition of pre-mRNA splicing substrates. Recent work uncovered an unexpected connection between mRNA splicing and cellular metabolism, in which genetic ablation of U1 snRNP components in erythroleukemia cells promoted a shift towards increased use of oxidative phosphorylation (OXPHOS), promoting growth in galactose-containing media (Jourdain et al. 2021). Of these components, knockout of *LUC7L2*, a non-constitutively bound U1 snRNP “auxiliary factor”, presented the strongest metabolic phenotype, resulted in altered splicing of several hundred exons. The shift toward OXPHOS could be partially explained by changes in the splicing of key metabolic genes, including *PFKM* and *SLC7A11*.

**Figure 1.**
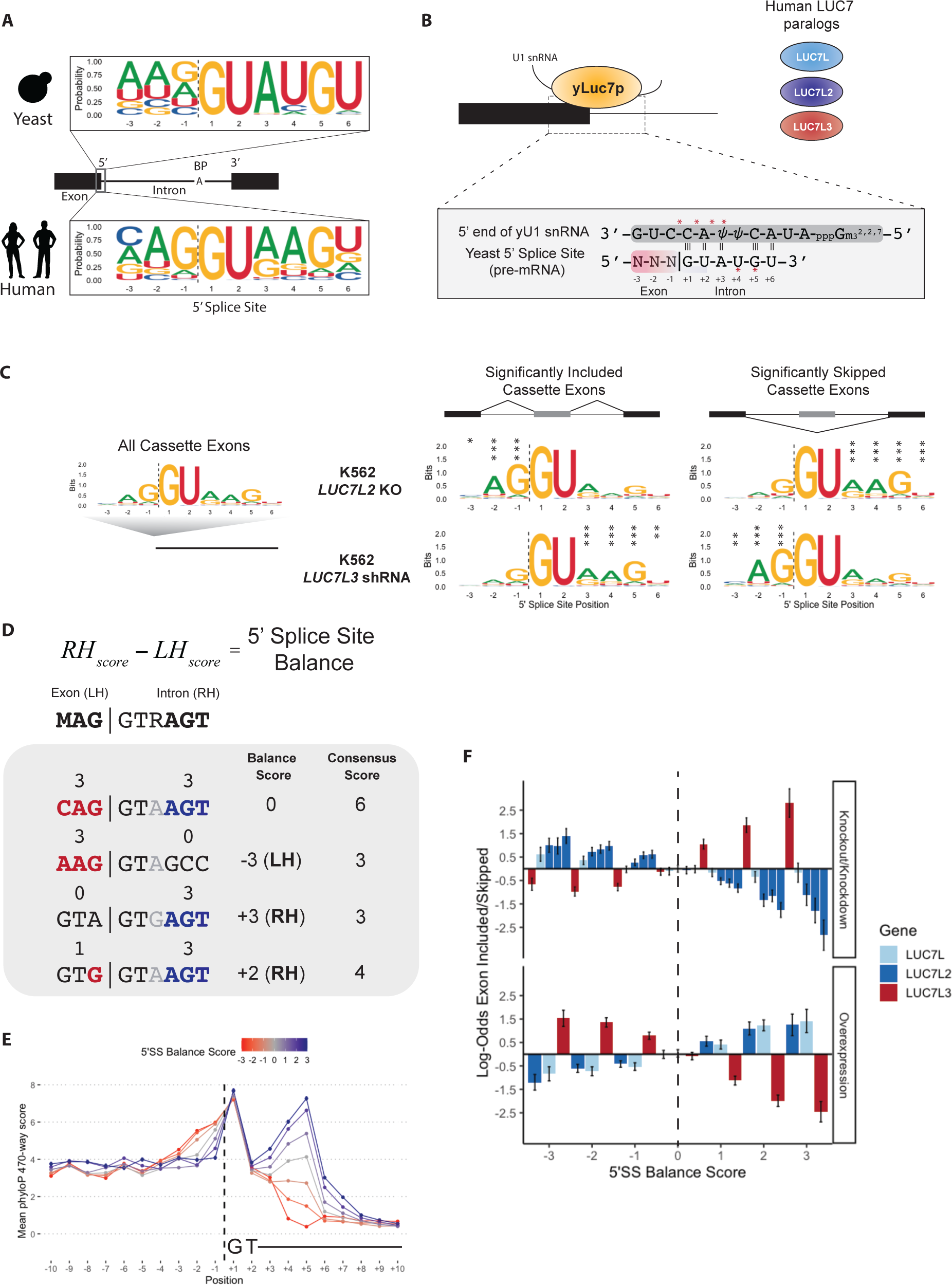
Human LUC7 proteins impact 5’ splice sites with distinct composition. A) Sequence logos of annotated 5’SS in yeast (top) and humans (bottom). B) Cartoon represen-tation of yLuc7p contacting U1 snRNA:5’SS duplex. Red asterisks indicate reported locations on duplex where yLuc7p interacts directly with U1 snRNA (Bai et al., 2018). C) Representative sequence logos of 5’SS for all cassette exons (left) and exons with significantly increased or decreased inclusion upon depletion of LUC7L2 or LUC7L3. D) Schematic of 5’SS Balance score, which is defined as the number of consensus bases at positions +4/+5/+6 minus the number of consensus bases at –3/–2/–1 of a 5’SS. E) Mean phyloP score for human 5’SS of each possible 5’SS Balance score. F) Log-odds of a 5’SS with a given 5’SS Balance score being differentially included versus skipped for each indicated human LUC7 RNA-seq dataset.

Previous work on the LUC7 family suggests a role in 5’SS selection (Fortes et al., 1999, Puig et al., 2007, Agarwal et al., 2016). Cryo-EM studies of the *S. cerevisiae* spliceosome showed that the yeast LUC7 is positioned adjacent to the U1 snRNA:5’SS duplex, interacting with the phosphodiester backbone of U1 snRNA during the initial stages of pre-spliceosome assembly (Bai et al., 2018, Plaschka et al., 2018, Li et al., 2019). While yeast possess a single, constitutively U1-associated LUC7 protein, yLuc7p, mammals possess three: the closely-related LUC7L and LUC7L2, and the more divergent LUC7L3 (Jourdain et al. 2021, Daniels et al. 2021). Immunoprecipitation of each human LUC7 protein recovered U1 snRNP proteins, confirming interaction with the snRNP. All three proteins crosslinked with splice sites in many pre-mRNAs and with bases at the 5’ end of U1 snRNA that basepair with the 5’SS (Daniels et al., 2021, Jourdain et al., 2021). Despite their high conservation to yLuc7p, their association with U1 snRNP, and broad impact on the transcriptome, human LUC7 proteins are absent from all published cryo-EM spliceosome structures.

Ablation of *LUC7L2*, but not of other LUC7 family members, promotes an oxidative phosphorylation metabolic state, supporting a functional distinction between these paralogs (Jourdain et al., 2021). In iPSC-derived hematopoietic stem cells, low expression and/or loss of *LUC7L2* via del(7q) impacts differentiation (Kotini et. al, 2015). Low expression and/or loss of *LUC7L2* is also associated with development of leukemia and other myleoid neoplasms (Singh et al., 2013), suggesting that LUC7L2’s influence on splicing and cellular metabolism may contribute to leukemogenesis. However, the molecular functions of the LUC7 paralogs remains incompletely understood.

Here, we investigated the distinct roles of LUC7s in pre-mRNA splicing. Unexpectedly, we found that different human LUC7 paralogs broadly impact the splicing of thousands of exons in a predictable, 5’SS-sequence-dependent manner. Experiments and analyses in a broad range of systems, from human cells to mutant plants, demonstrate that two subfamilies of LUC7 proteins regulate two newly identified classes of “left-handed” (LH) and “right-handed” (RH) 5’SS in opposing manners, helping to explain the distinct phenotypes of these paralogs.

## Results

### LUC7 family members impact distinct subclasses of 5’SS

In yeast pre-B complex, the second zinc-finger domain of yeast Luc7 (yLuc7p-ZnF2) forms salt bridges with the phosphodiester backbone of U1 snRNA while duplexed with the 5’SS (Bai et al., 2018) (Fig. 1B). The three human LUC7 family members each possess two N-terminal zinc finger (ZnF) domains, but LUC7L3-ZnF2 is highly divergent compared to its paralogs (Fig. S1A). Since human LUC7 paralogs also interact with human 5’SS (Puig et al., 2007, Jourdain et al., 2021, Daniels et al., 2021), we reasoned that the divergence of LUC7L/LUC7L2 and LUC7L3 may have resulted in distinct sets of binding/regulatory targets. Therefore, we examined the 5’SS features of dysregulated exons in publicly available RNA-seq data from knockout (KO) or knockdown (KD) of these genes in human cells, including *LUC7L2* KO in K562 erythroleukemia and HeLa cells, and *LUC7L*, *LUC7L2*, and *LUC7L3* KD in K562 cells. Analysis of 5’SS strength (Yeo and Burge, 2004) indicated that exons sensitive to depletion of LUC7 family members had either no significant difference in mean 5’SS strength relative to unchanged exons or had differences that were significant but extremely small in magnitude, ranging from 0 to 0.25 bits (Fig. S1B). These observations suggest 5’SS strength contributes at most minimally to exon responsiveness to Luc7 perturbations.

Despite having similar strength, the 5’SS of exons impacted by depletion of *LUC7L2* and *LUC7L3* had strikingly different sequence compositions compared to unchanged exons (Fig. 1C). Specifically, exons more skipped following *LUC7L2* KO had higher consensus matching at positions +3, +4, +5 and +6 on the intron side of the 5’SS motif, while those more included had increased consensus matching at positions –3, –2 and –1 in the exon. In contrast, KD of *LUC7L3* yielded an opposite pattern, with exons more skipped following depletion having higher consensus matching at –3, –2 and –1, and those more included having higher consensus matching at positions +3, +4, +5 and +6. No differences were observed in the sequence logos at the upstream 5’SS or at the upstream or downstream 3’SS relative to affected exons (Fig. S1C), supporting that the 5’SS sequence of the impacted cassette exon is important for LUC7 regulation.

Analysis of human 5’SS logos (Fig. 1C) suggests that the extent of consensus matching on each side of the /GU is an important determinant of sensitivity to *LUC7L2* versus *LUC7L3* depletion, and previous studies (Yeo and Burge, 2004; Carmel et al., 2004, Roca et al., 2008) have noted the preferential co-occurrence of adjacent consensus bases in the intron or exon portion of the 5’SS motif. To more quantitatively assess this feature, we devised a simple metric called the “5’SS Balance score” (or simply 5’SS Balance), which summarizes the distribution of consensus bases around the 5’SS. This score is calculated as the number of matches to consensus on the intron side at positions +4, +5 and +6 minus the number of consensus matches on the exon side at positions –3, –2 and –1 (Fig. 1D). At position –3, C or A are counted as consensus, and position +3 is ignored for the time being. 5’SS Balance ranges from –3 to +3, where LH 5’SS with more consensus matching on the exon side receive negative scores and RH 5’SS with more consensus matching on the intron side receive positive scores. “Balanced” exons with equal consensus matching on both sides of the /GU motif receive a score of 0.

Across all human exons, about 20% have Balance ≤ –2, while about 15% have Balance ≥ +2 (Fig. S2A). The 5’SS Balance score is a property distinct from 5’SS strength (with a weak negative correlation observed, Fig. S2B) and is generally conserved across vertebrates and between orthologous exons between human and mouse (Fig. 1E, Fig. S2C). Together, these results imply that LH and RH 5’SS subclasses are abundant and stably maintained as subclasses of 5’SS.

### 5’SS Balance predicts sensitivity to *LUC7L2* versus *LUC7L3*

To assess the extent to which 5’SS Balance could predict splicing changes in response to LUC7 depletion, we evaluated the log-odds of increased inclusion versus skipping across available LUC7 RNA-seq datasets. This analysis indicated that 5’SS Balance is strongly predictive, with LUC7L2 promoting the splicing of exons with RH 5’SS and repressing exons with LH 5’SS, and LUC7L3 promoting splicing of exons with LH 5’SS and repressing splicing of exons with RH 5’SS (Fig. 1F). Similar effects were observed for LUC7L2 KD in K562 cells and LUC7L2 KO in HeLa, while depletion of LUC7L gave results similar in direction to depletion of LUC7L2, but more modest in magnitude. To confirm these findings, we performed RNA-seq on HEK293T cells overexpressing each individual LUC7. Overexpression yielded the opposite effects on LH and RH exons as depletion for each paralog (Fig. 1F), supporting our conclusions that that splicing of LH 5’SS are promoted by LUC7L3, while RH 5’SS are promoted by LUC7L2 and LUC7L.

To systematically identify 5’SS motifs impacted by each LUC7 protein, we measured the impact of LUC7 proteins on the enrichment of each 5’SS 9mer (spanning positions –3 to +6) in exons differentially included or skipped. This measure, which we call “5’SS Enrichment”, uses a Dirichlet-multinomial model to approximate the log-odds of a given sequence occurring in significantly included or skipped versus unchanged exons in an experiment. Positive enrichment values indicate over-representation of a 5’SS sequence in included exons and negative values indicate enrichment in skipped exons, while values near zero indicate absence of bias for inclusion/skipping (or presence in unchanged exons only).

As a positive control, we evaluated our approach on a recently published RNA-seq data where mutant U1 snRNA (g.3a>c) was overexpressed (Shuai et al., 2019; Figure S3A). As expected, we recovered very strong and significant enrichment of 5’SS with +6G (complementary to U1 snRNA (g.3a>c) position 3) (Fig. S3B). 5’SS Enrichment analysis of LUC7 perturbation experiments yielded many 5’SS motifs with strong enrichment in each direction (examples shown in Fig. S3C). Hierarchical clustering of significant 5’SS Enrichment scores across all 8 LUC7 RNA-seq datasets yielded clear groups of 5’SS with many LH motifs promoted by LUC7L3 and repressed by LUC7L2 and LUC7L, and many RH motifs promoted by LUC7L2/LUC7L and repressed by LUC7L3 (Fig. S3D).

Our analyses indicate that we can empirically define LH and RH 5‘SS by their sensitivity to LUC7 proteins. Therefore, we defined a “LUC7 Score” as a ratio of position weight matrices for LUC7L2-promoted/LUC7L3-repressed RH motifs versus LUC7L3-promoted/LUC7L2-repressed LH motifs based on 5’SS Enrichment across seven LUC7 perturbations (Methods) (Fig. 2A). This score, which is strongly correlated with the 5’SS Balance (r = 0.86, p < 2.2e-16), predicted the direction of splicing change in a cross-validation test (Fig. 2B), performing slightly better than or similar to 5’SS Balance in every case. The LUC7 Score captures additional sequence features including the base at +3 and the relative importance of different non-consensus bases at each position. For example, LUC7L2-promoted 5’SS favor –1U while LUC7L3-promoted 5’SS favor –1A when not matching the consensus (G) at position –1. The LUC7 Score groups human 5’SS into two similarly-sized clusters (Fig. 2C). Comparison of our LUC7 Score to conventional 5’SS features shows that it is uncorrelated with splice site strength (*r*= 0.007, p = 0.65) and distinct from other previously studied features of 5’SS (Fig. 2D). The individual PWMs for LH and RH motifs are weakly, but significantly correlated with predicted minimum free energy of interaction with U5 and U6 snRNAs, respectively. Together, these findings indicate that the LUC7 Score can leverage sequence features from RNA-seq data to accurately classify exons whose splicing is dependent on different LUC7 family members.

**Figure 2.**
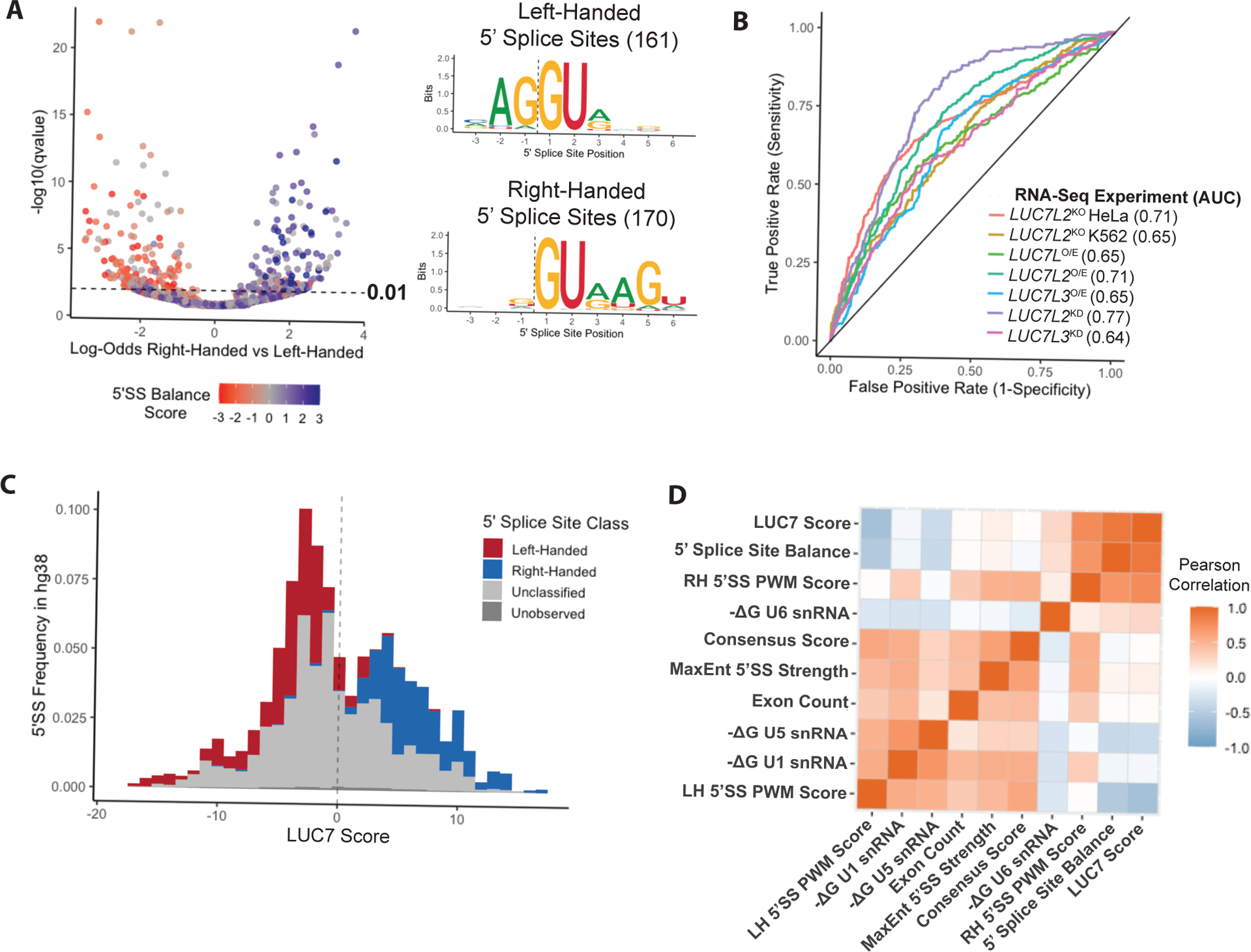
Enrichment of individual 5’ splice site motifs across LUC7 experiments defines two distinct classes. A) Volcano plot of 5’SS enrichment scores of individual 5’SS 9mers (each dot represents a 9-mer) from LUC7 meta-analysis (left). Sequence logos derived from significant 9mers (qvalue < 0.01) from LUC7 meta-analysis (right). B) Receiving operator characteristic curve of LUC7 score predictions of direction of regulation for held-out splicing events from LUC7 meta-analysis. C) Distribution of LUC7 score and frequency for distinct 9mer 5’SS sequences (with /GT) in Gencode human genome protein coding exons. D) Heat map showing Pearson correlation between new measures – 5’SS Balance and LUC7 score – versus standard 5’SS measures.

### LUC7 family members cross-regulate each other’s splicing via RH 5’SS

Many splicing regulatory factors (SRFs) negatively auto-regulate the expression of their own gene at the level of splicing, and are often negatively cross-regulated by close paralogs through the same exons (Spellman et al., 2007; Leclair et al., 2020). This regulatory arrangement, which helps to maintain a narrow range of activity for particular SRFs and SRF families, occurs in the LUC7 family as well (Daniels et al., 2021), with LUC7L2 repressing the expression of LUC7L by promoting the inclusion of an exon whose inclusion results in a premature termination codon (PTC) in LUC7L (Jourdain et al., 2021). Although such PTC-inducing exons are not annotated in the other LUC7 paralogs, we noted the strongly conserved alternative second exon (AE2) in *LUC7L2* that replaces the coding sequence of a highly conserved N-terminal alpha helix that interacts with SM-ring component SmE in U1 snRNP (Plaschka et al. 2018). Sequence alignment of the *LUC7L2* AE2 and *LUC7L* PTC exons revealed that they are 82% identical at the nucleotide level, implying a common evolutionary origin; no comparable exon was found in *LUC7L3*.

Consistent with previous studies, we found that overexpression of either *LUC7L* or *LUC7L2* promoted inclusion of the *LUC7L* PTC exon in a minigene assay (Fig. 3A), suggesting that both paralogs repress *LUC7L* expression (Fig. 3B,C, left). We found the *LUC7L2* AE2 is regulated in a similar manner to *LUC7L* PTC (Fig. 3B,C, right), supporting a pattern of negative auto- and cross-regulation analogous to that observed for *PTBP1*/*PTBP2* and other closely-related often partially-redundant SRF genes. In contrast, *LUC7L3* overexpression repressed the inclusion of the *LUC7L* and *LUC7L2* negative-regulatory exons, promoting their canonical mRNA isoforms.

**Figure 3.**
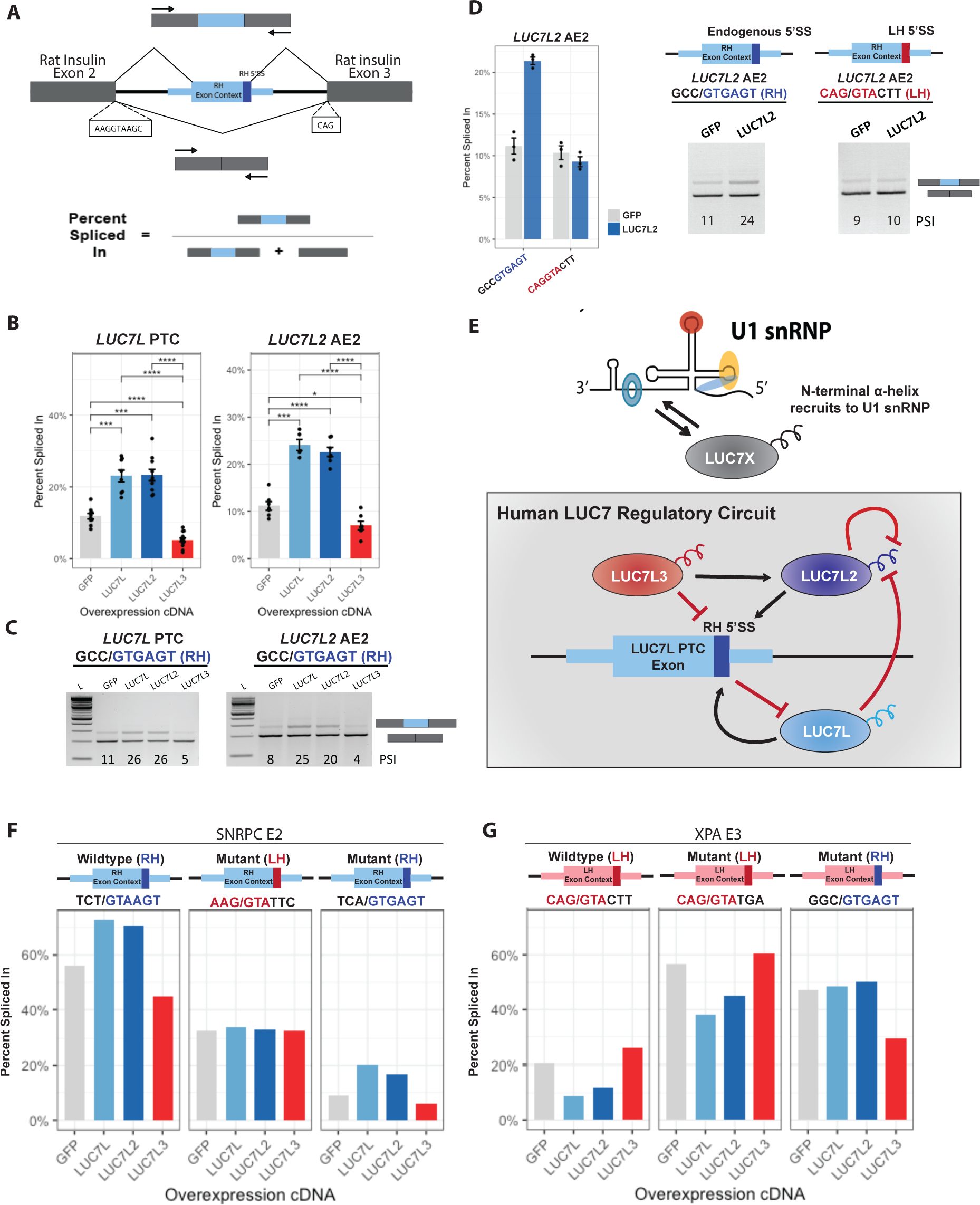
Regulation of splicing by LUC7 proteins depends on a 5’SS handedness. A) Schematic of pSpliceExpress minigene construct used, in which an internal exon of interest with flanking splice sites is inserted into a minigene expressing rat insulin exons 2 and 3 and intervening intron. B) Bar plots of cassette exon’s percent spliced in (by RT-PCR with primers in flanking exons) across all experiments; bars are color-coded to match LUC7 paralog colors in (D). Error bars represent standard error. Brackets indicate significant comparisons. *p < 0.05, **p < 0.01, ***p < 0.001, ****p < 0.00001, Bonferroni-corrected t-test. C) Representative gel images from LUC7-related minigenes. Percent Spliced In values shown at bottom of each lane. D) Mean PSI for LUC7L2 AE2 minigene with mutagenized 5’SS (left), and representative RT-PCR gel images (right). E) Proposed regulatory relationships between human LUC7 family members. F) Bar plot of PSI for SNRPC E2 minigene with wildtype RH 5’SS (left), mutagenized LH 5’SS (middle) and mutagenized RH 5’SS (right). G) Bar plot of PSI for XPA E3 minigene with wildtype LH 5’SS (left), mutagenized LH 5’SS (middle) and mutagenized RH 5’SS (right).

Notably, the 5’SS sequence of the homologous *LUC7L* PTC exon and *LUC7L2* AE2 are identical in sequence, of average strength and strongly RH in character (MaxEnt score = 8.7 bits, 5’SS Balance = +3 (RH), LUC7 Score = +7.6). To determine whether 5’SS properties contribute to regulation of these exons, we swapped the RH 5’SS of *LUC7L2* AE2 with a LH 5’SS of comparable strength (MaxEnt = 8.1 bits, Balance = –2, LUC7 Score = –8.4). We found that the autoregulatory *LUC7L2* AE2 exon with this new 5’SS was spliced similarly to the wildtype (WT) under control conditions, but that this change completely abolished splicing regulation by *LUC7L2* (Fig. 3D). Together, these observations indicate that the RH character of the AE2 5’SS is necessary for regulation by *LUC7L2*. Furthermore, it suggests human LUC7 proteins auto and cross-regulate one another through the 5’SS sequence of regulatory exons. Positive regulation by LUC7L3 may help to maintain a specific ratio of LUC7L2 + LUC7L to LUC7L3 activity, perhaps to ensure that both RH and LH 5’SS are spliced efficiently (Fig. 3E).

To assess the generality of the 5’SS-dependence of LUC7 regulation, we performed similar 5’SS mutagenesis of the RH 5’SS of *SNRPC* alternative exon 2, and of the LH 5’SS of *XPA* exon 3 in similar minigene contexts. As above, changing the *SNRPC* AE2 WT 5’SS (MaxEnt = 8.4 bits, Balance = +3, LUC7 Score = +13.7) to a LH 5’SS (MaxEnt = 7.6 bits, Balance = –3, LUC7 Score = –6.3) abolishes all positive and negative regulation by LUC7 proteins (Fig. 3F). However, mutation to a different RH 5’SS (MaxEnt = 8.3 bits, 5’SS Balance = +3, LUC7 Score = +9.7) preserved regulation similar to the WT sequence, demonstrating that RH 5’SS are necessary for positive and negative regulation by *LUC7L*/*LUC7L2* and *LUC7L3*, respectively. Similarly, switching the LH 5’SS of *XPA* exon 3 (MaxEnt = 8.1 bits, Balance = –2, LUC7 Score = –8.4) to a different LH 5’SS (MaxEnt = 9.5 bits, Balance = –2, LUC7 Score = –3.3) mimics the regulation of the WT LH 5’SS. However, splice site switching a LH 5’SS to a RH 5’SS (MaxEnt = 7.4 bits, Balance = +3, LUC7 Score = +10.6) abolished regulation by *LUC7L*/*LUC7L2* and yielded repression by *LUC7L3* (Fig. 3G).

These results indicate that RH 5’SS are necessary, but not sufficient, to confer positive regulation by *LUC7L* or *LUC7L2*, but may be sufficient to confer repression by *LUC7L3* (Fig. 3G). Altogether, our mutagenesis experiments suggest that the “handedness” (LH or RH character) of the 5’SS sequence is a key determinant of regulation by LUC7 family members.

### Specificity for RH 5’SS is seen by CLIP and conferred by LUC7 N-terminal domains

We hypothesized that positive regulation of RH 5’SS by *LUC7L2* occurs through a direct mechanism involving stabilization of U1 snRNP’s interactions with RH 5’SS by LUC7L2. To assess LUC7L2 RNA binding transcriptome-wide, we analyzed published eCLIP data (Jourdain et al. 2021). Consistent with previous analyses, LUC7L2 was found to crosslink predominantly to the 5’SS and a region ∼25 nt upstream of the 3’SS (Fig. 4A, top). To assess whether LUC7L2 crosslinks more to RH 5’SS, we binned exons by their LUC7 scores and then examined the Pearson correlation between the mean LUC7 score of the bin and eCLIP crosslinking enrichment in windows around the 3’SS and 5’SS of cassette exons. This analysis, performed on 4 eCLIP experiments in two separate human cell lines, revealed increased LUC7L2 crosslinking around the 5’SS of cassette exons with more RH 5’SS (Fig. 4A, bottom), supporting our hypothesis. Increased crosslinking was also observed to a region ∼25 nt upstream of the 3’SS near the expected location of U2 snRNP binding; this binding might result from cross-exon interactions of U1 and U2 snRNPs related to exon definition.

**Figure 4.**
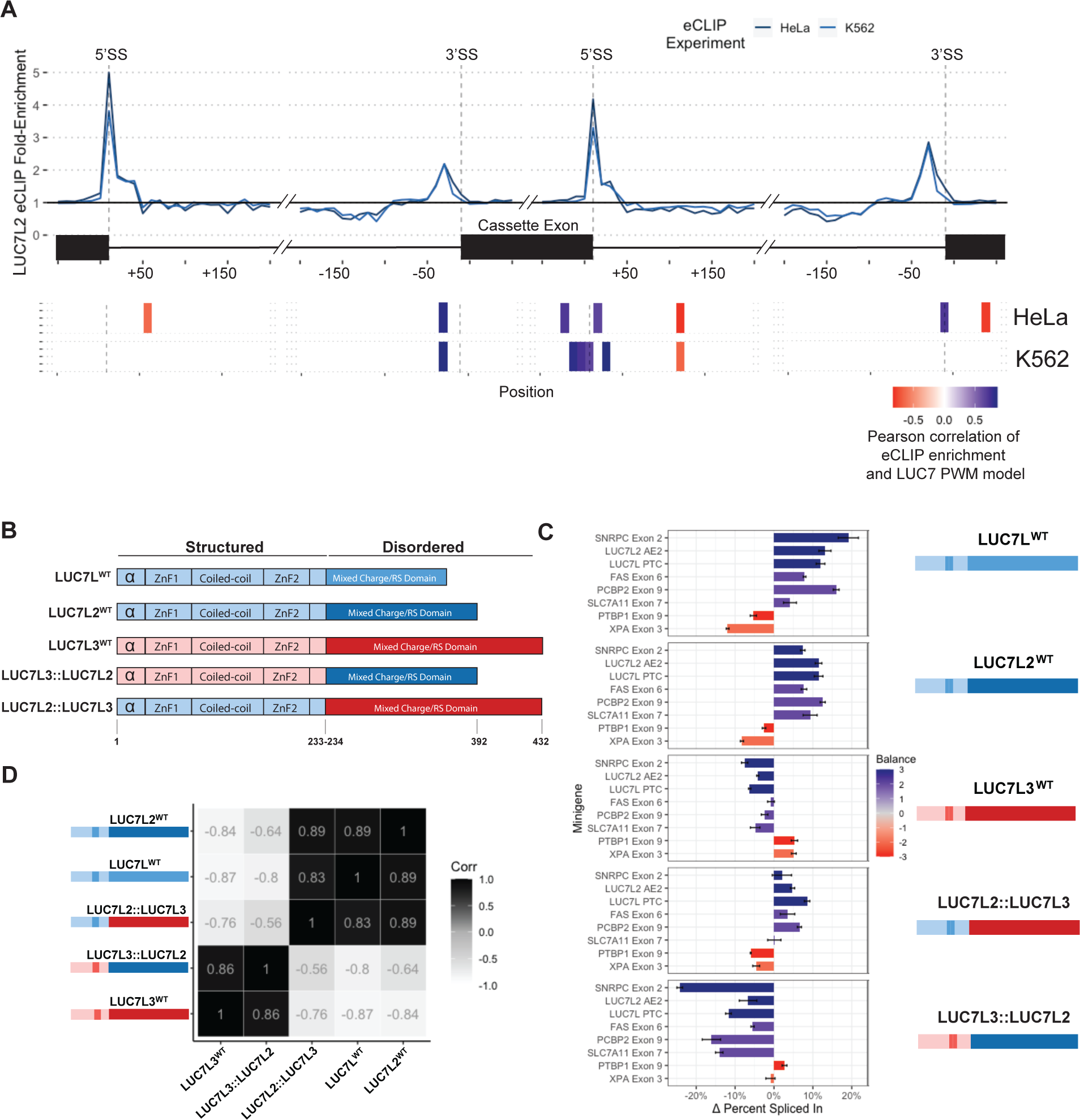
LUC7 specificity for subclasses of 5’SS depends on the structured N-terminal region. A) eCLIP enrichment of LUC7L2 around windows of constitutive and cassette exon splice sites. Crosslinks are aggregated into 10 nt bins (top). Pearson correlation of LUC7 score and LUC7L2 eCLIP enrichment, only significant (p < 0.05) correlations are reported. B) Protein domain structure of human LUC7 proteins and experimentally investigated chimeric proteins. C) Change (“delta”) in percent spliced in (qRT-PCR) from transfected minigenes containing different internal exons of introns following overexpression of LUC7 WT or chimeric cDNA (shown at right). Mean is plotted with error bars representing standard error, and bars are color-coded by the Balance score of the internal exon’s 5’SS. D) Heat map of Pearson correlations of delta percent spliced values for each LUC7 WT or chimeric cDNA overexpressed in C.

A multiple sequence alignment of human LUC7 paralogs with yLUC7p suggested that yLuc7p is more similar overall to LUC7L and LUC7L2, particularly in their ZnF2 domains (Fig. S1A), while LUC7L3 diverges in ZnF2, potentially contributing to functional differences relative to LUC7L/LUC7L2. Given that human LUC7L and LUC7L2 are more similar in sequence and possess similar target specificity in RNA-seq and minigene assays, we hypothesized that the divergent structured domains dictate LUC7 paralog specificity for distinct 5’SS subclasses. To test this hypothesis, we generated chimeric proteins by fusing the structured N-terminal regions of LUC7L2 or LUC7L3 with the C-terminal intrinsically disordered region (IDR) of the other paralog (Fig. 4B) and assessed the impact on exons with different types of 5’SS. Using a small library of minigene constructs with different RH and LH 5’SS exons of interest, over-expression of wildtype (WT) LUC7L or LUC7L2 invariably promoted the inclusion of exons with RH 5’SS and repressed inclusion of exons with LH 5’SS, while overexpression of WT LUC7L3 repressed inclusion of RH 5’SS and promoted inclusion of exons with LH 5’SS (Fig. 4C), consistent with the pattern established above. Considering the chimeric proteins, we found that LUC7L2 N-terminus fused to LUC7L3 C-terminus (LUC7L2::LUC7L3) yielded similar changes in splicing as LUC7L2 (or LUC7L) overexpression for both LH and RH 5’SS, though somewhat weaker in magnitude (Fig. 4C). On the other hand, the reverse chimera, LUC7L3::LUC7L2, repressed RH 5’SS inclusion, often somewhat more potently than WT LUC7L3, and activated one of the two LH 5’SS studied. These data and an associated correlation analysis (Fig. 4D) support that the structured N-terminal domains of LUC7L2 and LUC7L3 are sufficient to confer the general pattern of repression/activation of LH versus RH 5’SS, and suggest that the C-terminal regions of these paralogs have similar, but not identical functions in splicing.

### AMLs with low expression of *LUC7L2* have distinct splicing and expression patterns

Loss of one copy of *LUC7L2* occurs commonly with monosomy 7 or loss of the long arm of chromosome 7 (−7/del(q)), a cytogenetic feature common in both myelodysplastic disorders (MDS) and acute myeloid leukemias (AML) and invariably associated with poor survival (Gupta et. al, 2018). In a differentiation phenotype-rescue screen, *LUC7L2*, along with *EZH2*, *HIPK2* and *ATP6V0E2*, were identified as haploinsufficient genes responsible for mediating the hematopoietic defect associated with −7/del(q) (Kotinni et. al 2015). Furthermore, the *LUC7L2* locus is recurrently mutated in patients with MDS and AML, and low expression of *LUC7L2* was identified as a common characteristic among patients who harbor mutations in other MDS-associated splicing factors (Hershberger et. al, 2021). These links between *LUC7L2* and AML pathogenesis motivated us to focus on AMLs with variations in copy number (CNVs) for *LUC7L2* and to hypothesize that AMLs with reduced *LUC7L2* expression (LUC7L2^Low^) would have reduced inclusion of RH versus LH 5’SS compared to other AMLs. To test this hypothesis, we identified 14 AML samples harboring deletions spanning the *LUC7L2* locus. As expected, AMLs with fewer copies of *LUC7L2* express roughly 2-fold lower *LUC7L2* expression on average. While we did not detect differences in the expression of *LUC7L* or *LUC7L3* mRNA in these tumors, we observed that both the *LUC7L* and *LUC7L2* autoregulatory exons were less included in AML samples with low expression of *LUC7L2* (not shown), as predicted by our LUC7 regulatory model (Fig. 3E). Differential splicing analysis comparing 16 LUC7L2^Low^ AMLs versus 157 LUC7L2^Ctrl^ samples recapitulated our earlier results from LUC7L2-depleted human cell lines, in which differentially skipped and included exons possess right- and left-handed 5’SS sequences, respectively (Fig. 5C, Fig. S3E, S3F). Conducting the same splicing analysis on shuffled AML groups completely abolishes the observed relationship between 5’SS handedness and splicing regulation. Together, these analyses show that LUC7L2^Low^ AMLs inefficiently splice RH 5‘SS relative to LH 5’SS, supporting a role for reduced *LUC7L2* levels in shaping the transcriptomes of these tumors.

**Figure 5.**
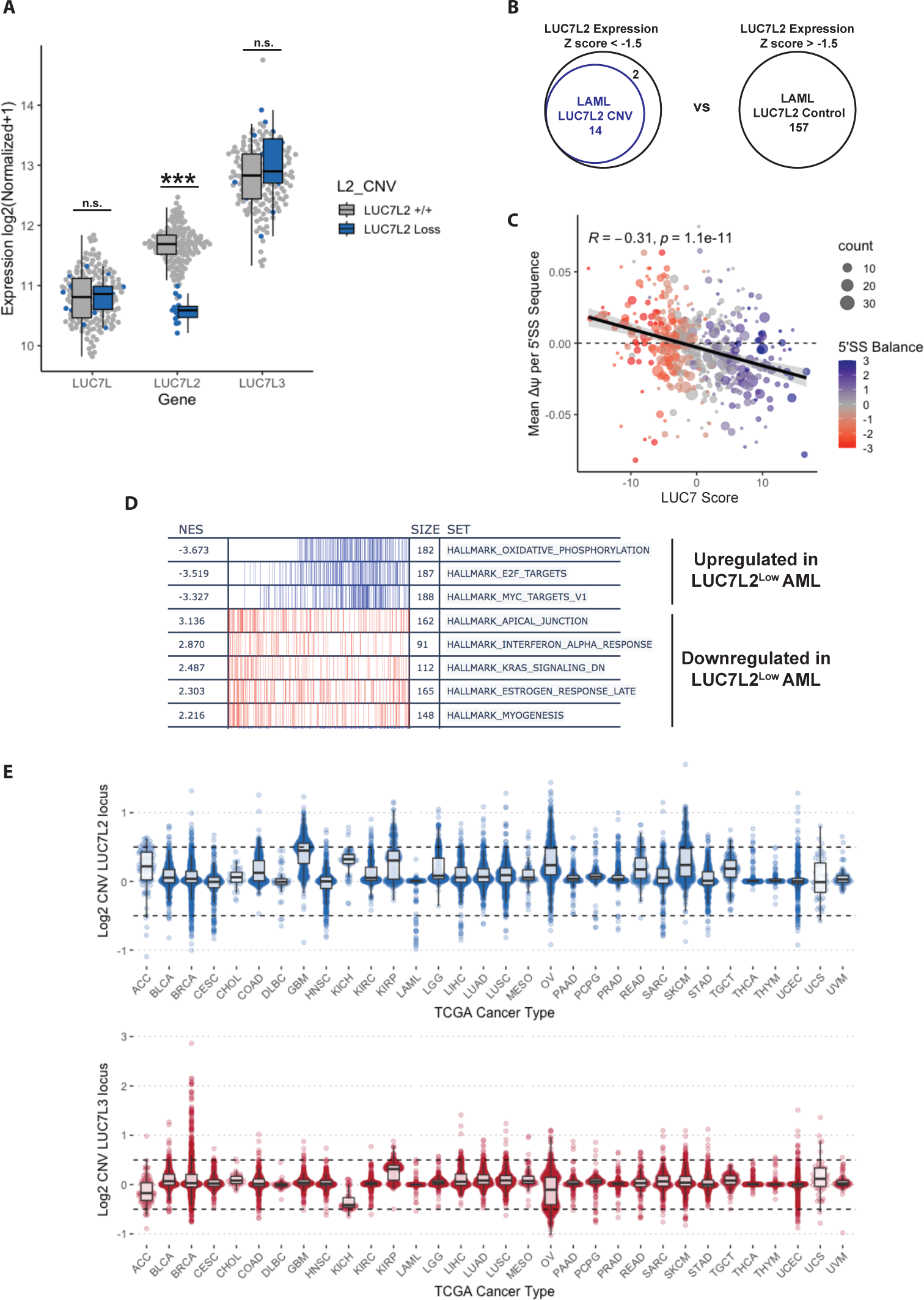
Transcriptomic analysis of Acute Myeloid Leukemias with low expression of LUC7L2. A) Normalized gene expression values for human LUC7 family in LAML samples, colored by whether samples possess a LUC7L2 CNV loss (log_2_ value < –0.5). B) Overlap of *LUC7L2* CNV loss samples and LUC7L2^Low^ expression samples. C) Mean dPSI per 5’ splice site sequence for differentially spliced exons when comparing LUC7L2^Low^ versus LUC7L2^Ctrl^ expression samples. D) Gene set enrichment analysis of differentially expressed genes comparing LUC7L2^Low^ versus LUC7L2^Ctrl^ expression samples. E) Copy-number variation analysis for *LUC7L2* and *LUC7L3* loci across all TCGA cancer types.

To assess potential functional consequences of low LUC7L2 expression in AML, we performed gene set enrichment analysis (Fig. 5D). These analyses revealed that LUC7L2^Low^ AMLs have increased expression of genes associated oxidative phosphorylation relative to other AMLs, suggesting that perturbation of LUC7L2 expression in AMLs may mimic the metabolic state observed in K562 *LUC7L2^KO^* cells (Jourdain et. al., 2021). Other pathways associated with cell cycle and DNA damage repair, such as E2F target genes, are also generally increased. Altogether, these results signify 5’SS subclasses are differentially regulated by LUC7 proteins in AML and that low expression of LUC7L2 may promote a transcriptomic and/or metabolic state conducive to AML pathogenesis (de Beauchamp et al. 2022).

To explore whether variation of LUC7L2 and LUC7L3 may be involved in other cancers, we examined the CNV of these genes across the entire TCGA cohort (CNV data for *LUC7L* were not available). This analysis revealed that several tumor types, including ACC, GBM, KIRP, OV, SKCM and TGC exhibit increased copy number of *LUC7L2*, with some of these (ACC, KICH, OV) showing decreased copy number of *LUC7L3*, suggesting dysregulation of U1 snRNP and potential skewing of 5’SS selection toward RH 5’SS in these cancers (Fig. 5E). Additionally, some cancer types (e.g., BRCA, KIRP, UCS) had increased copy number of *LUC7L3*. Some of these observations reflect well-established chromosomal aberrations commonly found in specific tumor subtypes, e.g., trisomy 7 is common in glioblastomas (GBM), yielding an additional copy of *LUC7L2*. Changes in LUC7 expression due to CNVs may generally contribute to the observed splicing variation found in many cancer subtypes and might contribute to metabolic changes as well.

### Evolutionarily-related Luc7 proteins possess conserved specificities for 5’SS subclasses

To understand the origins of the divergence in function between the *LUC7L3* and *LUC7L2/LUC7L*, we searched for homologs of these proteins and performed multiple sequence alignments. We identified at least one annotated Luc7 family member in 7 out of 9 major eukaryotic supergroups (Fig. S5A), indicating that this gene family is extremely ancient. We also performed multiple sequence alignment of Luc7 proteins in 33 plants, animals and fungi, spanning a deep split in eukaryotic phylogeny (Burki et al., 2020). The resulting dendrogram supports that there are two major subfamilies of Luc7 proteins, represented by LUC7L2 and LUC7L3, and that both of these subfamilies were present before the split between plants and animals/fungi (Fig. 6A). Human LUC7L2 and LUC7L, which have similar 5’SS specificity via our minigene and transcriptome analyses, duplicated more recently in the last common ancestor (LCA) of jawed vertebrates approximately 425 million years ago (Fig. 6B). Intriguingly, while animals and plants retained both LUC7L2-like and LUC7L3-like proteins, none of the analyzed fungi possess a member of the LUC7L3 subfamily (see below).

**Figure 6.**
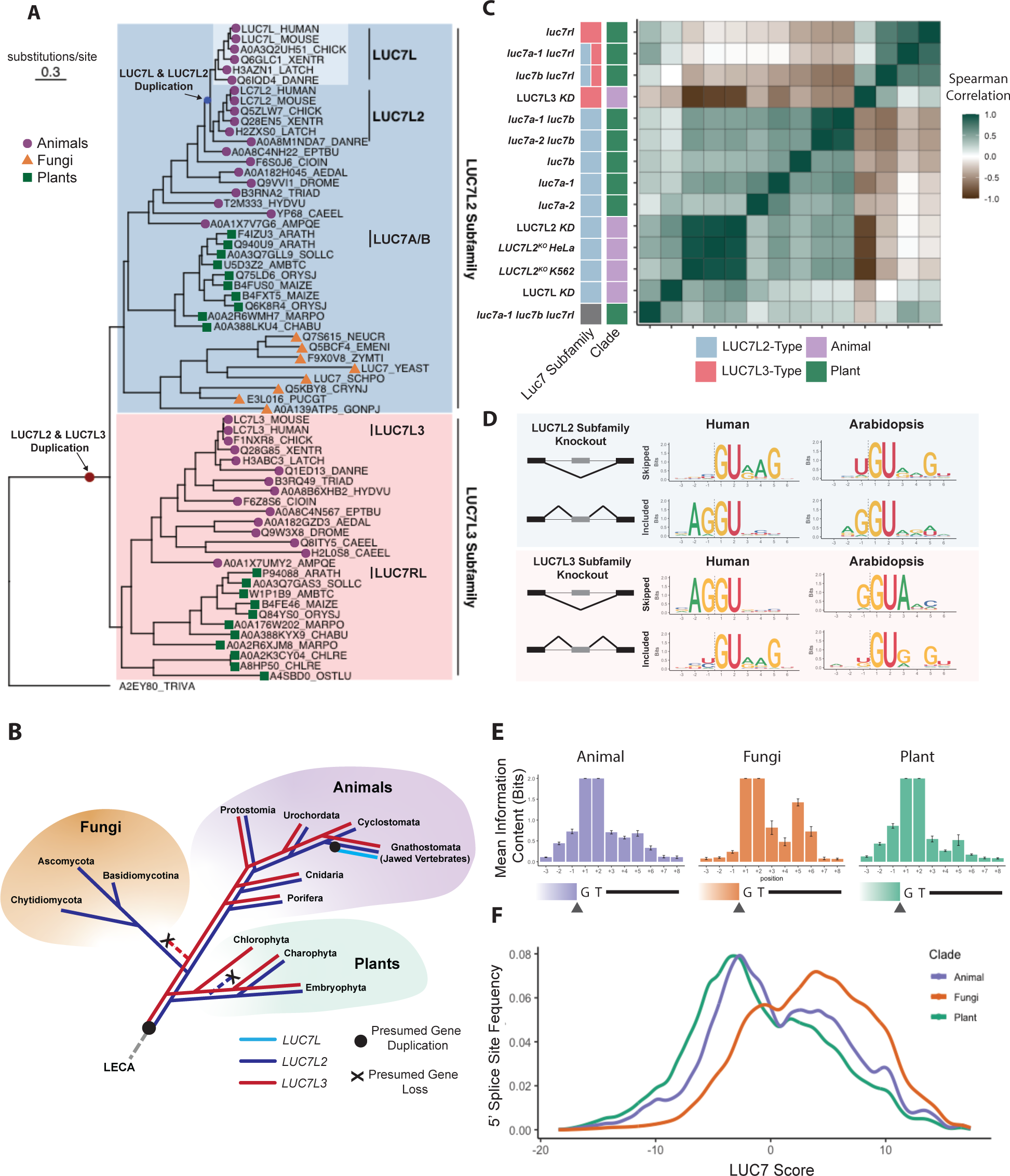
Phylogenetic analysis suggests coevolution of Luc7 proteins and 5’SS motifs. A) Maximum likelihood phylogenetic tree built from multiple sequence alignment of Luc7 proteins from 33 animal, fungal and plant species, adding *Trichomonas vaginalis* Luc7 protein as an outgroup. Two main clusters are shaded to indicate Luc7 subfamilies; individual proteins represented by symbols indicating clade of origin. B) Representation of presence/absence and likely duplication/loss events for Luc7 subfamilies, overlaid on the eukaryotic phylogenetic tree. C) Correlation matrix of 5’SS models learned from dinucleotide features of 5’SS differentially spliced in human and plant Luc7 RNA-seq experiments. D) Sequence logos of top or bottom 10% unique 5’ splice sequences identified by dinucleotide 5’SS models. E) Mean position-specific information content (calculated as in Irimia et al., 2019) of 5’SS motifs color-coded by clade. F) Density plot reflecting mean 5’SS subclass frequency in each eukaryotic clade studied.

The deep conservation of Luc7 family members across animals and plants provides an opportunity to compare the molecular functions of Luc7 proteins across eukaryotic supergroups. *Arabidopsis thaliana* has three Luc7 genes: two members of the LUC7L2-like subfamily (*LUC7A* and *LUC7B*) and one member of the LUC7L3 subfamily (*LUC7RL*), which are important for development and stress resistance (de Francisco Amorim et al., 2018). To assess whether evolutionarily-related LUC7 proteins possess analogous 5’SS specificity, we performed RNA-seq on every possible combination of *Arabidopsis thaliana* luc7 single, double and triple mutants, and carried out differential splicing analysis.

Consistent with previous work, analysis of cassette exons revealed that luc7 single mutants have relatively modest splicing phenotypes: *luc7a* and *luc7b* showed a trend toward higher LUC7 score in skipped exons (like *LUC7L2*), while luc7rl showed the opposite trend (like *LUC7L3*), but only luc7a reached statistical significance (Fig. S5B-C). Strikingly, a similar analysis of the *luc7a/luc7b* double mutant showed a very strong bias for RH 5’SS among skipped exons, similar to depletion of *LUC7L2*. This observation held true for *luc7a/luc7b* double mutants with two different alleles of *luc7a* (Fig. S5D). Similarly, *luc7rl* mutants trended toward negative LUC7 scores among skipped exons, reaching significance in one case (Fig. S5D). No bias for LH or RH 5’SS in differentially skipped or included exons was observed in the luc7 triple mutant.

To assess the overall similarity of the 5’SS dysregulated in human and plant LUC7 datasets, we devised a model which identifies 5’SS nucleotide pairs (consecutive or non-consecutive) as predictive features of 5’SS dysregulated within an RNA-seq data set. Scoring all human 5’SS with each model, and assessing the correlation between these scores, we identified two major clusters: one containing *LUC7L3*, *luc7rl* and *luc7rl*-containing double mutants; and the other containing LUC7L2, LUC7L and all of the *A. thaliana* mutants involving *luc7a* and/or *luc7b* (Fig. 6C). Furthermore, as predicted by our dinucleotide model, the top 10% 5’SS sequences most sensitive to LUC7 subfamily depletion are very similar between human and plants. In both human and *Arabidopsis*, depletion of LUC7L2 subfamily members consistently results in more skipping of RH 5’SS with a +5G and −1U, while depletion of LUC7L3 subfamily members results in more skipping of LH 5’SS with both –1G and +5H (Fig. 6D). These observations indicate that the two subfamilies of LUC7 proteins in plants have distinct activities on 5’SS subclasses, paralleling our observations in humans.

While the 5’SS subclasses impacted by human and plant LUC7 proteins are very similar overall, we do observe some species-specific differences. For example, *Arabidopsis* LUC7RL promotes LH 5’SS with −1G, rather than LH 5’SS with a –2A/–1G pair promoted by LUC7L3 in humans. These subtle differences may reflect lineage-specific preferences for particular 5’SS substrates by distantly-related LUC7 proteins. Together, these results support that orthologous human and *A. thaliana* LUC7 proteins retained their ancestral specificities for their respective 5’SS subclasses over 1.5 billion years of evolution.

### Loss of the *LUC7L3* subfamily is associated with altered 5’SS composition

Our observations that LUC7L3-subfamily genes promote LH 5’SS and LUC7L2-subfamily genes promote RH 5’SS in both *A. thaliana* and human imply that the 5’SS specificity of these two subfamilies evolved prior to the divergence of animals/fungi from plants. Since none of the fungal species studied possess a LUC7L3 ortholog, it appears likely that this subfamily was lost from the fungal lineage soon after its divergence from animals, so that fungal genomes have evolved largely in the absence of a LUC7L3. To explore patterns of 5’SS composition, we compiled annotated /GT 5’SS from the genomes of 33 eukaryotes, representing plants, animals and fungi, and summarized the positional information content in each lineage (Fig. 6E). This analysis revealed distinct patterns of information content between the three clades, with greater similarity between organisms within each clade. Consistent with previous studies, all analyzed fungi have strongly biased composition on the intron side of the 5’SS, with little or no bias at any exonic position (Kupfer et al., 2004). Meanwhile, all animals, except for 3 Protostomes (insects and nematodes) studied, have moderate biases at both exonic and intronic positions (Lim and Burge, 2001). Plant 5’SS have slightly stronger bias on the exon side than animals, but are otherwise similar to animals than either is to fungi (Szczesniak et al., 2013).

Assessing 5’SS subclasses, we observed that while plants and animals (other than Protostomes) have roughly comparable frequencies of LH versus RH 5’SS, the RH class of 5’SS is predominant in fungi (Fig. 6F). Together, these data suggest long-term coevolution between LUC7 subfamilies and 5’SS subclasses, with loss of the LUC7L3 subfamily and depletion of LH 5’SS both occurring early in the evolution of fungi.

## Discussion

In both animal and plant lineages, our findings support a model where LUC7L2/LUC7L and LUC7L3 play significant roles in the regulation of splicing by tuning the recognition of distinct subclasses of 5’SS. We propose that LUC7L2 and LUC7L3 interact with U1 snRNP in a manner that differentially promotes or stabilizes binding to right-handed (RH) and left-handed (LH) 5’SS, respectively. Structural evidence suggests that LUC7 proteins directly interact with the U1:5’SS duplex (Bai et al., 2018), while biochemical assays demonstrate LUC7 proteins associate with U1 snRNP and crosslink near the 5’SS. Notably, LUC7L2 crosslinking near 5’SS correlates with the right-handedness of the splice site, emphasizing specificity for different 5’SS subclasses. The mechanism underlying negative regulation of the opposite 5’SS subclass could involve destabilization, or could reflect competition of different LUC7 family members for access to the same site on U1 snRNP.

Additional evidence consistent with our model comes from recent work in *S. cerevisiae* where humanized LUC7L/LUC7L2-ZNF2 rescued cell viability and promoted more flexible 5’SS recognition, contrasting with the inviability of yeast complemented with humanized LUC7L3-ZNF2 (Carrocci et. al, 2024). These findings are consistent with our own experimental and evolutionary observations, in which LUC7L and LUC7L2 are functionally redundant, promote splicing of exons with fungal-like RH 5‘SS, and the activity of human LUC7L3 is distinct from its paralogs.

While we favor a differential U1 snRNP stabilization model, an impact on later stages of spliceosome assembly is possible. For example, yLUC7p is found as late as pre-B complex, the stage at which the tri-snRNP is recruited to splicing substrates. In the catalytic spliceosome U5 and U6 can basepair with exonic and intronic positions of the 5’SS, repsectively (Lesser and Guthrie, 1993; Newman and Norman, 1992, Sontheimer and Steitz, 1992). An activity of LUC7s that promoted or stabilized one of these interactions could potentially favor use of LH or RH 5’SS, respectively. Further biochemical and structural studies are needed to elucidate the mechanisms by which animal and plant LUC7 proteins act.

LUC7 proteins are unlikely to be the only *trans*-acting factors that impact the splicing of 5’SS subclasses and other studies in model organisms *A. thaliana* and *S. pombe* highlight the involvement of U5 and U6 snRNAs. U6 snRNA is subject to an S-adenosylmethionine-dependent m6A modification in the ACAGAGA box, which base pairs with an intronic portion of 5’SS. Recent research has shown that METTL16-mediated m6A modification promotes the use of 5’SS which have a RH character, highlighting the complexity 5’SS choice (Ishigami et al., 2021, Parker et al., 2022).

Our finding linking specific LUC7 proteins with specific subclasses of 5’SS may have significant implications for the future advancement of small molecule regulators of splicing. The synthetic lethal relationship between *LUC7L* and *LUC7L2* suggests potential therapeutic avenues for AML patients with monosomy 7 or other splicing factor mutations, who may benefit from therapeutics that specifically interfere with recognition of RH 5’SS. Similarly, if human METTL16 also impacts RH 5’SS recognition through U6 snRNA modifications, inhibition of METTL16 or methionine metabolism (Cunningham et. al, 2022) may be more effective in AML patients with low expression of *LUC7L2*.

## Methods

### Cloning

Human LUC7L, LUC7L2 and LUC7L3 ORFs were amplified from human cDNA and cloned into pcDNA3.1(+)IRES GFP (Addgene #: 51406). Domain swap constructs were synthesized as gblocks from IDT and cloned into pcDNA3.1(+)IRES GFP. Exons and flanking intronic regions used for pSpliceExpress minigenes were PCR amplified from human male genomic DNA using primers with attB overhangs and subsequently recombined into pSpliceExpress using BP Clonase II (Thermo Fisher, cat no. 11789020)

### Cell Culture & Chemical Transfection

HEK293T RMCE cell lines were cultured in Advanced DMEM supplemented with 5% FBS, 25 mM HEPES and Glutamax. For each experiment, cells were plated 24 hours in advance in 24-well plates. The following day, cells were transfected with 500 ng of 95:5 w/v of a cDNA overexpression vector and minigene reporter respectively using Lipofectamine LTX (Thermo Fisher, cat. no. 15338100)

### Minigene Reporter Assay

RNA was extracted 24 hours after transfection using Qiagen RNeasy Mini kit (cat. no. 74104) according to manufacturer’s instructions with the optional on-column DNAse digestion (cat. No 79254). RNA was eluted in nuclease free water and quantified using Nanodrop. For each RNA sample, we used 125 ng of RNA input into a 12.5 uL LunaScript Multiplex One Step Master Mix for RT-PCR (cat. no. E1555S). After PCR cycling, we mixed PCR samples with NEB 6X loading dye and loaded 5 uL of PCR products on a 3% agarose gel infused with ethidium bromide, which was run for 40 min at 150V. Except for Figures 3F and 3G, all presented minigene data was performed at least in triplicate. Images were acquired using Azure Biosystems c600 with UV imager.

### Percent Spliced In Calculation

Agarose gel images were manually quantified using chromatograms in ImageJ. Percent Spliced In values were calculated by taking the signal intensity of the larger band and dividing it by the sum of the signal intensity of the included product and the skipped product.

### LUC7 Overexpression and RNA-seq

Flag-tagged human *LUC7L*, *LUC7L2* and *LUC7L3* ORFs were transfected into HEK293 RMCE cells as described above. RNA was extracted 24 hours after transfection using Qiagen RNeasy Mini kit (cat. no. 74104) according to manufacturer’s instructions with the optional on-column DNAse digestion (cat. No 79254). RNA was eluted in nuclease-free water and quantified using Nanodrop. Illumina-compatible libraries were prepared by MIT BioMicroCenter using NEB II Ultra Directional RNA with poly(A) selection and sequenced on NovaSeq 6000 with 2 x 150 bp reads.

### Analysis of LUC7 paralog RNAseq data

We downloaded RNAseq data for *LUC7L2* KO in HeLa and K562 erythroleukemia cells (GSE157917) and *LUC7L*, *LUC7L2*, *LUC7L3* KD in K562 cells (E-MTAB-9709). For splicing analyses, we used rMATS 4.1.1 (Shen et al., 2014) with default settings.

In any given RNA-seq experiment, only a subset of exons likely change in splicing, with most other exons having Δψ near zero. The Δψ estimates reported by rMATS, however, are direct summaries of the read counts and are affected by sampling variation in the read counts which may artificially inflate changes for exons with low read counts. Commonly, shrinkage estimates are used in differential expression analyses to account for this issue. Shrinkage considers the set of all effect sizes to constrain the noise in estimates from low read count events. Doing so, however, requires parameter estimates and associated uncertainty, which are not directly reported by rMATS.

For each exon, we reconstruct the effect size in log-odds scale δ from the rMATS read counts and approximate a standard deviation σ describing the uncertainty in δ using the rMATS p-value. We pass these effect sizes and standard deviations to ashr using the ‘normal’ option, which assume that the proportions of up- and down-regulated exons are equal (Stephens, 2017). Using these shrinkage estimates we reconstruct an estimate of Δψ^∗^ (see Supplemental Methods).

### 5’SS Enrichment Calculation

We used a Dirichlet-multinomial model to calculate the log-odds of whether a given 5’SS was more likely to be involved in a significantly included event vs significantly skipped event. For each dataset, we excluded all events with fewer than 10 junction-count reads on average across all samples. First, we combined the counts of 5’SS from significant included and skipped events (FDR < 0.1) and their respective background sets, which consisted of an equal number of unregulated 5’SS that were matched for both PSI and expression level, into a count matrix. To avoid division by zero, we added a pseudocount of 1 to every observed 5’SS sequence and also accounted for class imbalance by dividing each column by a “weight” that reflected the fraction of included or skipped events. For example, if there were 4,000 significantly included events and 1,000 significantly skipped events, the significantly included counts and its respective background set was divided by 0.8 and skipped events and its respective background was divided by 0.2.

Then, we used this count matrix as the alpha parameters for Dirichlet-multinomial model, and simulated drawing from the posterior distribution 2500 times. For each draw we calculated the log-odds of a given 5’SS being enriched in the included versus skipped set. The posterior distribution of log-odds generated from the Dirichlet-multinomial model was used to calculate the posterior mean and the posterior standard deviation, which were both passed to ashr for shrinkage using the uniform option. From the output, we plotted the PosteriorMean estimates in 5’SS Enrichment plots, which can be directly interpreted as the log-odds of a given sequence occurring in the differentially included exon set over the differentially skipped exon set. For hierarchical clustering, we used scaled 5’SS enrichment scores to cluster 5’SS sequences by their activity across each LUC7 paralog RNA-seq dataset using Euclidean distance and ward.D2 linkage.

### 5’SS Enrichment meta-analysis

To calculate a single statistic describing the likelihood of a given 5’SS to be LH (LUC7L2-repressed/LUC7L3-promoted) versus RH (LUC7L2-promoted/LUC7L3-repressed) using all of the LUC7 RNA-seq datasets (except for LUC7L KD), we performed a modified 5’SS enrichment analysis. For each dataset, we applied the same filtering criteria as above and used a significance threshold of FDR < 0.1 to group exons in differentially-included, differentially-skipped or background sets. We then set aside a random 12.5% fraction of differentially included or skipped events as the held-out test set (Fig. 2B) to evaluate our LUC7 score. Background sets of unregulated exons for differentially included or skipped exon sets were matched for PSI and expression and were threefold larger in size. For each of these 4 groups, we sampled 5,000 exons with replacement. This approach was repeated for every RNA-seq dataset (except for *LUC7L* knockdown from Daniels et al., 2021). The events were then aggregated into a single table such that each RNA-seq data was equally represented in the final 5’SS enrichment analysis.

To account for opposing effects on LH and RH 5’SS subclasses for different experiments, the direction of differentially included and skipped events (and their respective background sets) from *LUC7L3* KD, *LUC7L* OE and *LUC7L2* OE analyses were flipped, such that included events represented RH 5’SS and skipped events represented LH 5’SS. Then we performed a 5’SS enrichment analysis as described above using a Dirichlet-multinomial model and simulated drawing from the posterior 10,000 times. As above, the distribution of log-odds generated from the Dirichlet-multinomial model was used to calculate the posterior mean and the posterior standard deviation, which were both passed to ashr for shrinkage using the uniform option. From this output, we plotted the PosteriorMean estimates in 5’SS enrichment plots and the associated qvalue. LH 5’SS (LUC7L2-repressed/LUC7L3-promoted) were defined as 5’SS with negative meta-5’SS enrichment scores with qvalue < 0.01 and and RH 5’SS (LUC7L2-promoted/LUC7L3-repressed) were defined as 5’SS with positive meta-5’SS enrichment scores with qvalue < 0.01.

### Position Weight Matrix and LUC7 score calculation

Sequence logos were created from LH and RH 5’SS sequences identified from meta-5’SS enrichment analyses. Individual PWM were created by calculating the observed frequency of a nucleotide at a given position, assuming a uniform nucleotide distribution. Pseudocounts of 0.1 were used to avoid division by zero. The LUC7 score was calculated by taking the ratio of the LUC7L2-promoted/LUC7L3-repressed RH PWM over the LUC7L3-promoted/LUC7L2-repressed LH PWM.

### Free energy predictions of U1, U5, U6 and 5’SS sequences

We used ViennaRNA 2.5 (Lorenz et al., 2011) to model free energy predictions between all 5’SS and the 5’end of U1 snRNA (ATACTTACCUG), U6 snRNA (ATACAGAGA) and U5 snRNA loop 1 (GCCUUUUAC) using RNAcofold with default parameters.

### LUC7L2 eCLIP analysis

Publicly available LUC7L2 eCLIP data was downloaded from European Read Archive (PRJNA663333). 10 nucleotide UMIs were extracted from reads and appended to read name. Then illumina adaptors were removed from reads using cut-adapt with the following settings (-a AGATCGGAAGAGCACACGTCTGAACTCCAGTCA \ --minimum-length 18 \ --quality-cutoff 6 \ --match-read-wildcards \ -e 0.1). This was performed twice to ensure complete removal of adaptor sequences. Using STAR 2.7.3a, we created a genomic index using GRCh38.primary_assembly.genome.fa and gencode.v38.primary_assembly.annotation.gtf with --sjdbOverhang 65. Trimmed eCLIP reads were then aligned to this genome using STAR with default settings and resulting bam files were then deduplicated with umi-tools. Crosslink counting was restricted to cassette exons observed in LUC7L2 KO datasets around 250 nt windows surrounding four splice sites (upstream 5’SS, cassette exon 3’SS, cassette exon 5’SS and downstream 3’SS) and crosslinks in 10 nt bins were summed together.

For both cell lines, 2 pulldowns and 2 size-matched (SM) inputs were considered. Crosslink counts in each library were first normalized by library size. Cassette exons were then stratified into 10 bins using their 5’SS LUC7 score (1 = most LH, 10 = most RH) and the number of crosslinks for each LUC7 score bin were summed together. Position-based eCLIP enrichment was calculated as the ratio of normalized eCLIP crosslinks to normalized SM-input crosslinks at each 10 nt bin. eCLIP enrichment within each 10 nt bin was then correlated with the mean LUC7 score of binned cassette exons.

### Phylogenetic and cross-kingdom splice site analysis

For our phylogenetic analysis, we manually chose 13 animals, 9 fungi and 11 plants from Esembl that represented diverse species in the each of the eukaryotic kingdoms. For each of these species, we used BLAST from UniProt to identify putative Luc7 paralogs and performed multiple sequence alignment using MAFFT (Katoh & Standley, 2013). We then created a maximum likelihood tree with RAxML (Stamakis, 2006) and annotated the final protein tree using ggtree (Yu et al., 2016). For each of the species used in our phylogenetic analysis, we downloaded their genomes and associated gene annotation files from Ensembl and extracted their splice site sequences using bedtools 2.29.2. For each species, we filtered for only /GT splice sites and calculated the information content per position of the 5’SS.

### *Arabidopsis thaliana* lines and growth conditions

All *A. thaliana* mutant lines used in this study are in Columbia (Col-0) background and were previously described (de Francisco Amorim et. al, 2018). Seeds of Wildtype Col0, luc7 single mutants (luc7a-1, luc7a-2, luc7b-1 and luc7rl-1), luc7 double mutants (luc7a-1 luc7b-1, luc7a-2 luc7b-1, luc7a-1 luc7rl-1 and luc7b-1 luc7rl-1) and a luc7 triple mutant (luc7a-2 luc7b-1 luc7rl-1) were surface sterilized with chlorine-gas and then grown on half-strength Murashige Skoog (MS) plates containing 0.8% phytoagar in continuous light at 22°C for 10 days. Seedlings were collected and flash frozen in liquid nitrogen. Total RNA was isolated using RNeasy® Plant Mini Kit (Quiagen, cat. nos. 74904) according to the manufacturer’s instructions. mRNA stranded library preparation and sequencing (PE150) was done by Novogene (Cambridge, United Kingdom) using an Illumina Novaseq6000 system.

### Patient stratification for TCGA analysis and differential splicing analysis

TCGA patient GDC-PANCAN.masked_cnv and normalized RNA-seq expression values were downloaded from https://xenabrowser.net/datapages/?cohort=GDC%20Pan-Cancer%20(PANCAN)&removeHub= https://xena.treehouse.gi.ucsc.edu%3A443). To assess per-sample CNV for LUC7 loci, we filtered for genomic segments spanning *LUC7L2* (chr7:139341360-139422599) and *LUC7L3* (chr17:50719544-50756213) for each patient sample. AML samples with average log2 CNV < −0.5 were considered to have *LUC7L2* loss. To distinguish *LUC7L2* low-expressing AMLs from AML samples expressing control levels of LUC7L2, we converted the normalized gene expression for LUC7L2 to Z scores and selected 16 patients with Z score < –1.5 were selected as LUC7L2 low expressing AMLs. Of the 151 LAML with RNA-seq data, 14 LUC7L2 low samples were available through GDC Data Portal. Differential splicing analyses using rMATs was performed to compare these LUC7L2 low expression with the remainder of LAML cohort. As a control, we shuffled the patient labels and carried out the same differential splicing analysis.

### GSEA of LUC7L2^Low^ vs LUC7L2^Ctrl^ AML samples

Gene set enrichment was performed in the Xena browser web portal. The same 16 samples identified as LUC7L2 low expressing AMLs were selected and compared to the remainder of the cohort with available data.

### Dinucleotide 5’SS model and scoring

To identify and score dinucleotide features that contribute to a 5’SS to be significantly included or skipped, we calculated the frequency in which a given adjacent and non-adjacent dinucleotide pairs were found in the 5’SS of a differentially spliced exon. We took the observed dinucleotide frequency and divided it by a background frequency, calculated from a sample of an equal number of exons from the background or “unregulated” set. Similar to our 5’SS enrichment score, this value reflects the log-odds of a dinucleotide feature appearing in differentially included exons versus differentially skipped exons. To estimate the variance around this statistic for each dinucleotide pair, we sampled an equal number of exons from the unregulated set 100 times for all observed pairs of dinucleotides. We then passed the mean and standard deviation of these distributions to ashr for shrinkage using the uniform option. The shrunken mean of significant dinucleotide 5’SS features (qvalue < 0.01) for each RNA-seq dataset were retained and used for subsequent scoring. To score a 5’SS, the dinucleotides present in the 9mer were scored, and the posterior log-odds associated with each were summed.

## Acknowledgements

We thank the staff of the MIT BioMicro Center for Illumina NovaSeq library preparation and sequencing. We thank members of the Burge lab, as well as David Bartel, Nima Jaberi-Lashkari, Alexis Jourdain, Byron Lee, Vamsi Mootha, Phillip Sharp, Yigong Shi, and Gordon Simpson for their helpful discussions. This work was supported by grant GM085319 from the NIH (C.B.B.) and DFG grant LA2633-4/2 (S.L.).

## Author Contributions

C.J.K designed, performed and analyzed all experiments and computational analyses under supervision of C.B.B. A.C performed mouse and human 5’SS analyses under supervision of C.J.K. M.P.M. assisted with the development of statistical approaches and eCLIP analyses. S.S. and S.L. performed Arabidopsis RNA-seq experiments. C.J.K. and C.B.B. wrote the manuscript with input from all authors. All authors contributed to edits/revisions of the manuscript.

## Supplementary Information

Figures S1 to S5

