## Supplementary Figures for "LUC7 proteins define two major classes of 5’ splice sites in animals and plants"

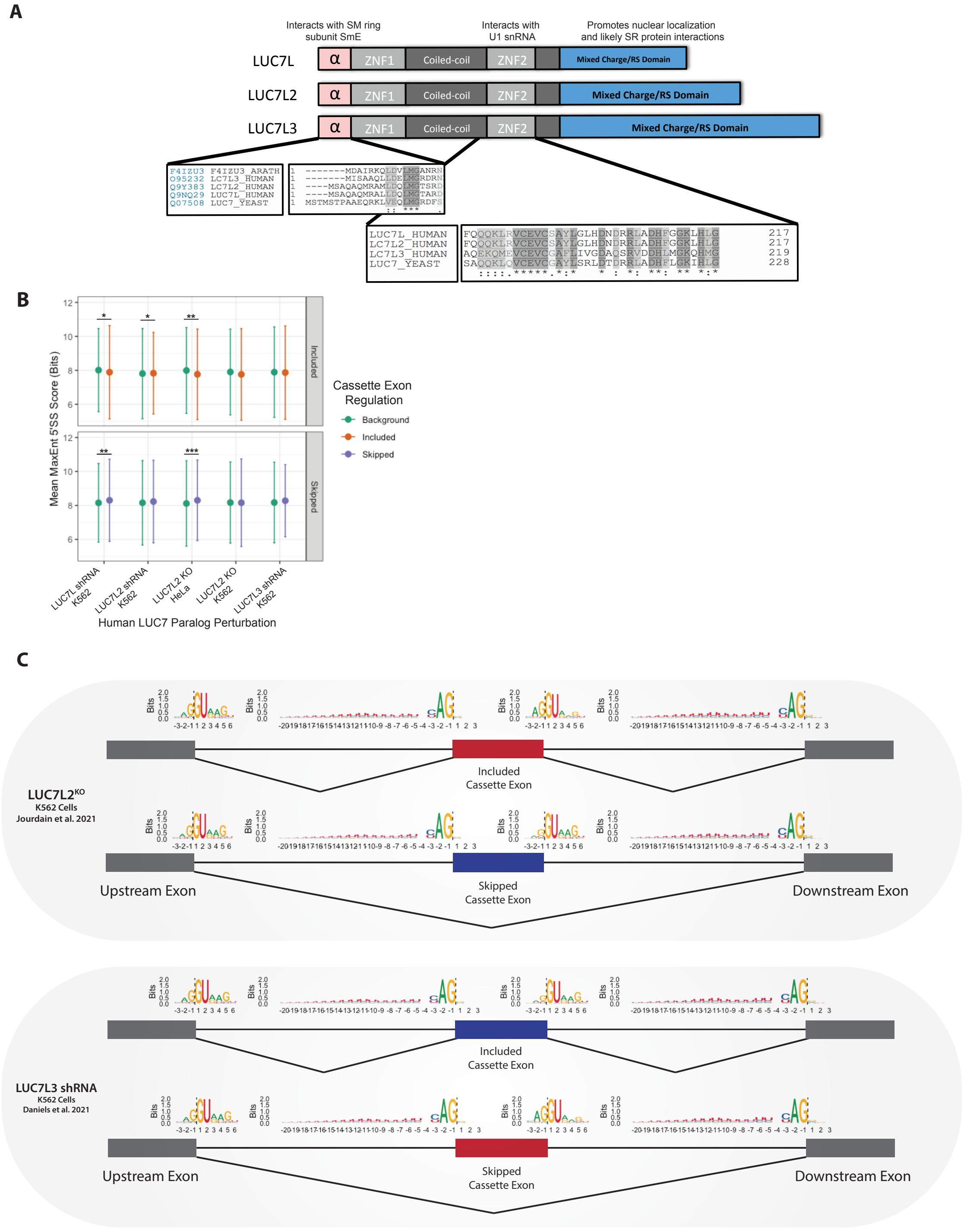


**Figure S1. 5' splice site motifs, strength and LUC7 paralogs.**
A) General domain structure of human LUC7 proteins (top) with multiple sequence alignment of N-terminal alpha helix that interacts with SM ring subunit SmE and ZNF2 (bottom). B) Mean 5'SS strength and standard deviation of LUC7-regulated exons and their respective background sets of unchanged exons matched for starting expression and PSI value. *p < 0.05, **p < 0.01, and ***p < 0.001, Wilcoxon rank sum test relative to respective background set. C) Summary of splice site sequence logos at upstream and downstream exons of LUC7-regulated cassette exons.

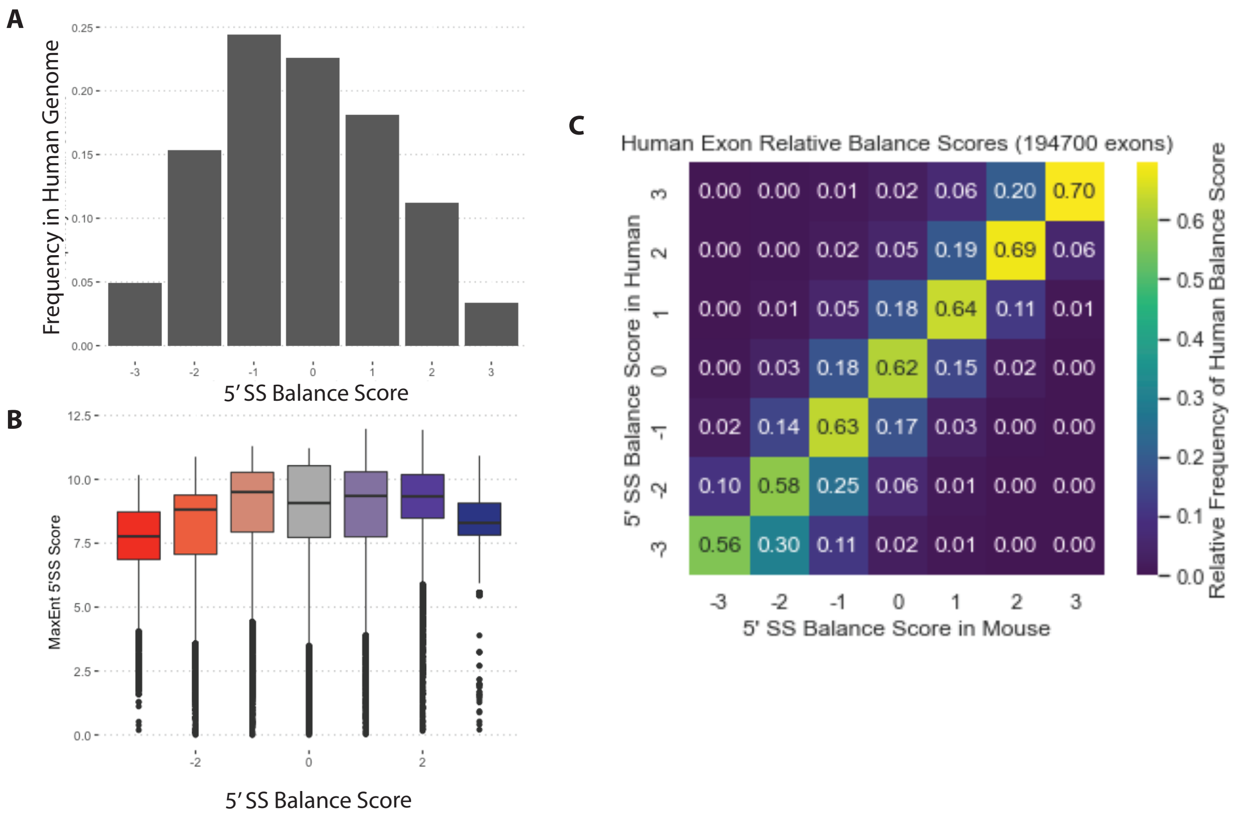


**Figure S2. 5'SS Balance scores of human and mouse orthologous exons.**
A) Frequency of 5'SS across 5'SS Balance scores in human protein-coding genes. B) Relationship of 5'SS strength and 5'SS Balance score in human protein-coding genes. C) Heatmap reflecting conservation of 5’SS Balance scores in orthologous mouse and human exons. Values in cells reflect proportion of conservation relative to human 5'SS Balance score. For example, 0.70 or 70% of 5'SS with +3 Balance score in humans have +3 5'SS Balance score in mouse.

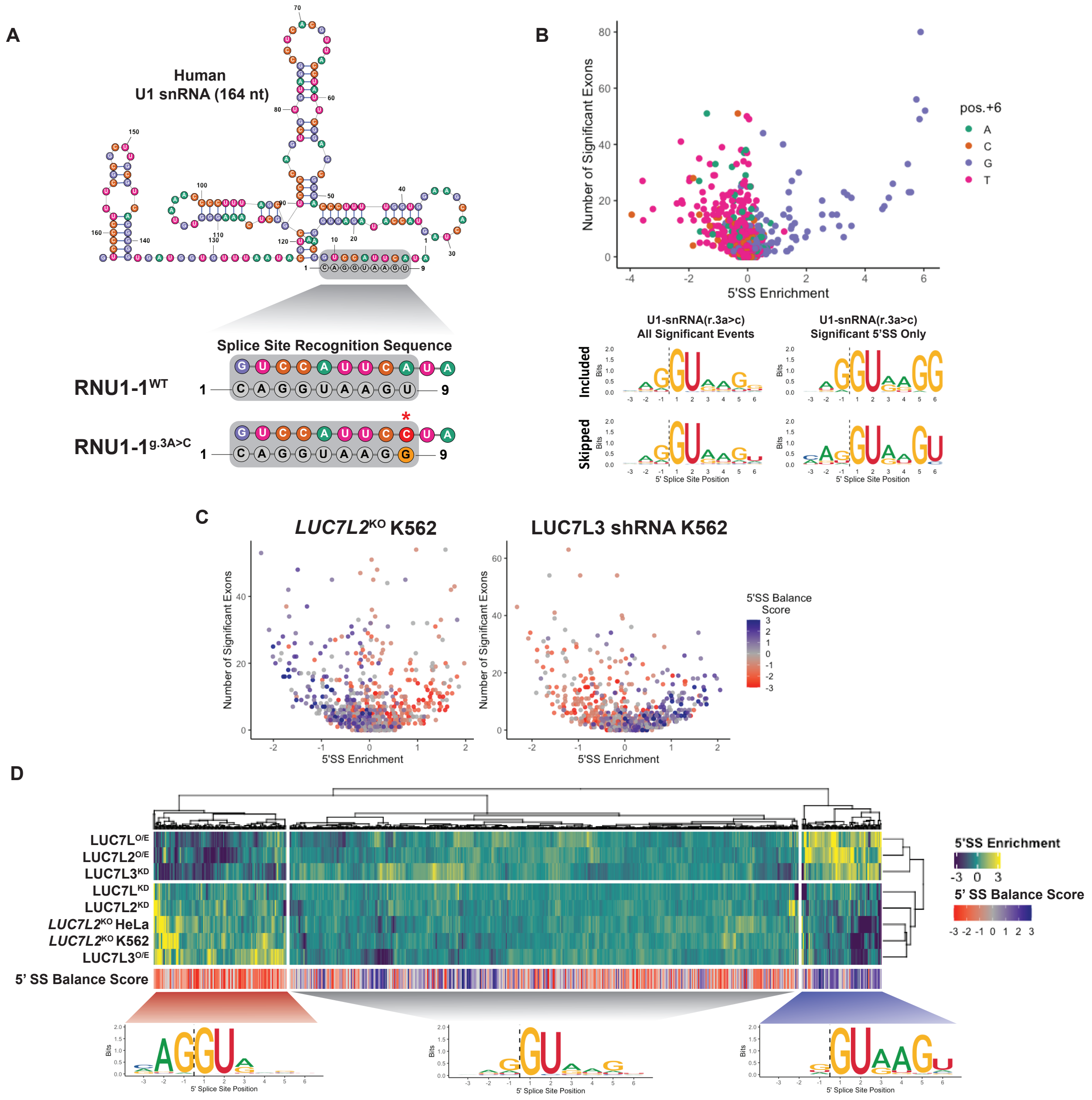


**Figure S3. 5'SS Enrichment application example and LUC7 depletion analysis.**
A) Simplified schematic of 5' end of WT (top) and mutant (bottom) U1 snRNA 5' end contacting the 5'SS. B) 5'SS enrichment scores of significantly changed cassette exons from overexpression of mutant U1 snRNA, colored by base at position +6. Sequence logos of 5'SS from all significant cassette exons (left) and from individual 5'SS with significant ($q$-value ≤ 0.01) 5'SS Enrichment values (right). C) Volcano plots of 5'SS enrichment analyses for LUC7L2 KO and LUC7L3 KD. D) Heat map of 5'SS Enrichment scores across all LUC7 RNA-seq datasets, clustered using Euclidean distance and ward.D2 linkage.

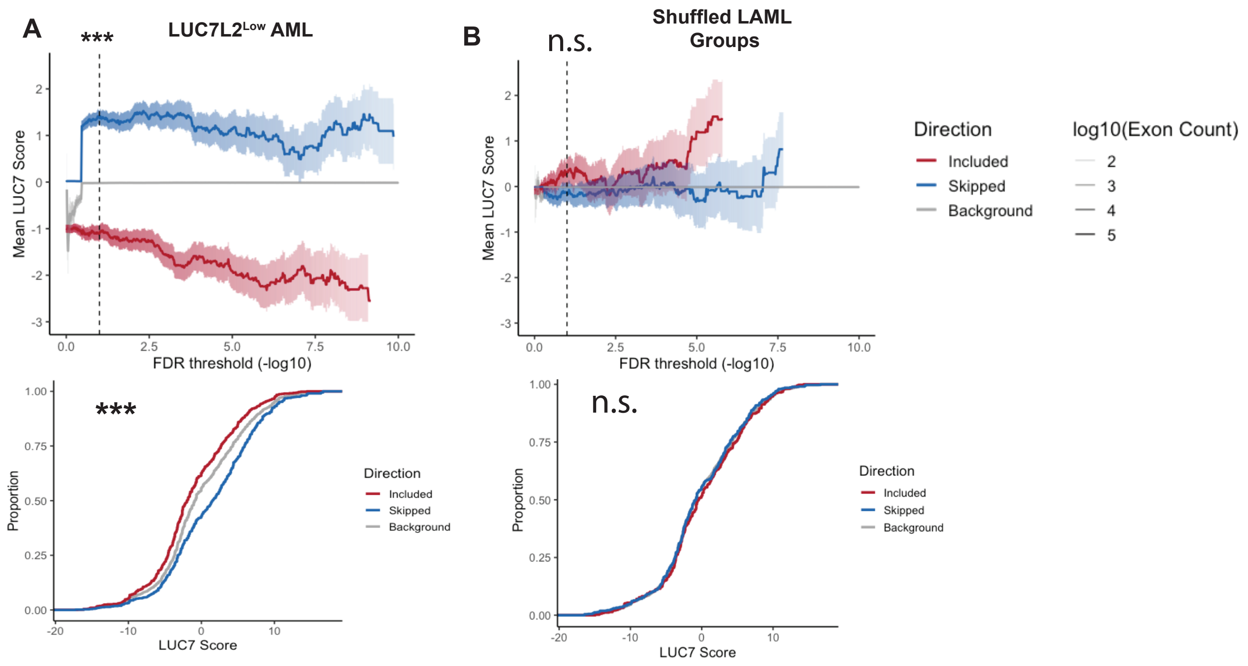


**Figure S4. Additional feature analysis of differentially regulated exons with acute myeloid leukemias with low expression of *LUC7L2*.**
A) FDR plot of LUC7 score for LUC7L2$^{Low}$ versus LUC7L2$^{Ctrl}$ samples, with CDF plot of LUC7 scores for exons at chosen FDR threshold. Dashed line indicates chosen FDR threshold of 0.1 for statistical test. B) FDR plot of LUC7 score for shuffled LAML samples, with CDF plot of LUC7 scores for exons at chosen FDR threshold. Dashed line indicates chosen FDR threshold of 0.1 for statistical test. *$p < 0.05$, **$p < 0.01$, ***$p < 0.001$, Wilcoxon-rank sum test.

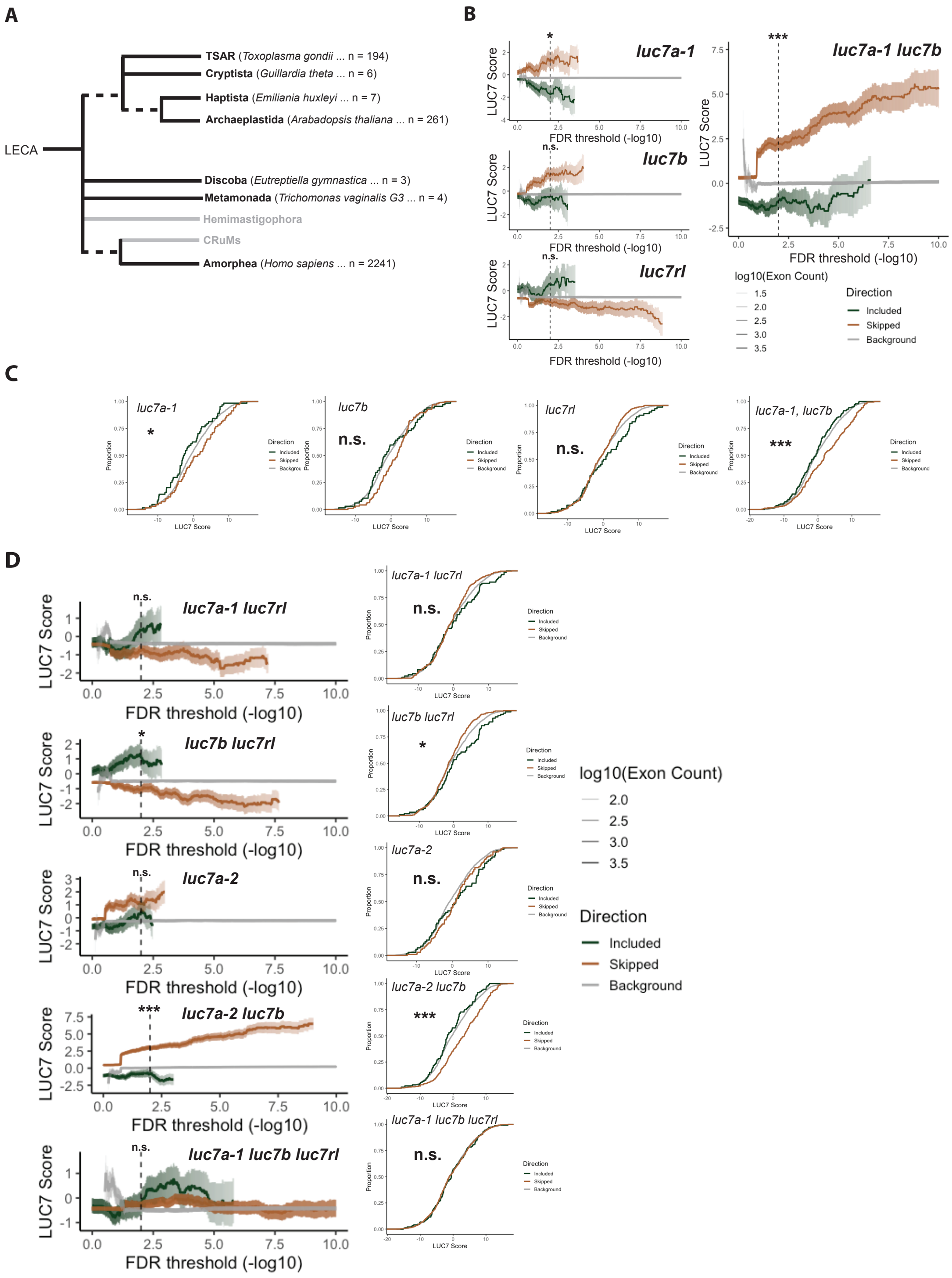


**Figure S5. Analysis of Luc7 genes and splicing phenotypes in *Arabidopsis*.**
A) Cladogram of eukaryotic supergroups summarized by Berki et al., 2020. Black supergroups contain an annotated LUC7 protein. An example species is provided next to each supergroup name, followed by the number of species with a LUC7 gene. Grey supergroups currently lack an annotated LUC7 protein. B) Relationship of mean LUC7 score as a function of FDR-threshold for single knockout *A. thailana* luc7 single mutants and one luc7 double mutant: *luc7a-1 luc7b*. Solid line is the mean of the examined feature and shaded regions indicate standard errors. Dashed

line indicates FDR threshold of 0.01 for Wilcoxon rank-sum test to compare features of significantly included vs significantly skipped events. A minimum of 50 exons are required for each group to be plotted. C) eCDF plots of luc7 single mutants and luc7 double mutant presented in (A). D) Additional FDR plots for all other Arabidopsis mutants. Dashed line indicates chosen FDR threshold of 0.01 for statistical test. *p < 0.05, **p < 0.01, ***p < 0.001, Wilcoxon-rank sum test.
